# Reprogramming of fish somatic cells for nuclear transfer is primed by *Xenopus* egg extract

**DOI:** 10.1101/2022.08.25.505281

**Authors:** Nathalie Chênais, Aurelie Le Cam, Brigitte Guillet, Jean-Jacques Lareyre, Catherine Labbé

**Affiliations:** INRAE, UR1037 LPGP, Fish Physiology and Genomics, Campus de Beaulieu, F-35000 Rennes, France; Université de Rennes 1, Campus de Beaulieu, F-35000 Rennes, France

## Abstract

Somatic cell reprogramming *in vitro* prior to nuclear transfer is one strategy expected to improve clone survival during development. In this study, we investigated the reprogramming extent of fish fin somatic cells after *in vitro* exposure to *Xenopus* egg extract and subsequent culture. Using a cDNA microarray approach, we observed drastic changes in the gene expression profile of the treated cells. Several actors of the TGFβ and Wnt/β-catenin signaling pathways, as well as some mesenchymal markers, were inhibited in treated cells, while several epithelial markers were upregulated. This was associated with morphological changes of the cells in culture, suggesting that egg extract drove somatic cells towards a mesenchymal-epithelial transition (MET), the hallmark of somatic reprogramming in induced pluripotent stem cells (iPSCs). However, treated cells were also characterized by a strong decrease in *de novo* lipid biosynthesis metabolism, the lack of re-expression of *pou2* and *nanog* pluripotency markers, and absence of DNA methylation remodeling of their promoter region. In all, this study showed that *Xenopus* egg extract treatment initiated an *in vitro* reprogramming of fin somatic cells in culture. Although not thorough, the induced changes have primed the somatic chromatin for a better embryonic reprogramming upon nuclear transfer.

## INTRODUCTION

In fish, somatic cells and particularly fin cells are a convenient source of diploid material for cryopreservation of valuable genetic resources ^[1]^. Such use of somatic cells compensates for the impossibility to cryopreserve fish oocytes and embryos. Besides, fin cells are easy to collect whatever the sex, maturation status or size of the fish, and they are easy to cryopreserve ^[2,3]^. However, regeneration of fish from these highly differentiated cells requires to master nuclear transfer, a technology that is still not reliable enough in fish. Indeed, whereas nuclear transfer with embryonic donor cells yields acceptable development rates ^[4–8]^, only few clones were reported to reach adulthood when the donor cell was taken from adult fish ^[9–16]^. One hypothesis often proposed to explain the low success rate of somatic cell nuclear transfer is the chromatin reprogramming failure (reviewed in mammals ^[17]^). In fish, zebrafish clones at dome stage fail to re-express several genes that are important for chromatin remodeling, translation initiation or cell cycle^[18]^. More recently, we showed that DNA methylation of several marker genes in goldfish clones failed to match the hypomethylation status of control embryos, and some clones bore the hypermethylated pattern of the donor fin cells ^[19]^. This means that after nuclear transfer, exposure of the somatic chromatin to oocyte factors prior to embryonic genome activation is not sufficient to overcome somatic cell resistance to reprogramming in fish.

Numerous studies in mammals have sought to improve the reprograming ability of donor somatic cells by way of an *in vitro* pre-reprogramming before nuclear transfer. The ability of metaphase-II (MII) egg factors to ensure chromatin remodeling of sperm and oocyte chromatin following fertilization makes the egg extract an attractive candidate for *in vitro* reprogramming. Heterologous *Xenopus* eggs at MII stage have been reported to improve the blastocyst rates after nuclear transfer in mouse ^[20]^, ovine ^[21]^ and porcine ^[22,23]^, and although assessed in only few studies, it also increased the development success after implantation or birth ^[21]^. At the molecular level, these heterologous egg extracts have also been shown to induce transcriptional and epigenetic remodeling in mammalian cultured cells. However, such reprogramming of somatic cultured cells is not straightforward and it suffers high variability. Many factors such as the animal species and cell type ^[24,25]^, the culture conditions and egg extract batches or stages ^[26,25,27,20]^ influenced the extent of somatic cell reprogramming. For example, when considering the expression of pluripotency markers, porcine cells treated with *Xenopus* egg extract only transiently re-expressed *Oct4* over culture time ^[26,28]^ and *Nanog* expression failed to be consistently re-expressed ^[26,29]^ while in mouse, *Oct4* and *Nanog* were both re-expressed ^[30,20]^. Moreover, the extent of reprogramming at the scale of the whole genome is not known. Indeed, all studies are assessing the reprogramming success from candidate genes analysis, and a reprogramming assessment based on all other putative actors of pluripotency is still missing. As a consequence, knowledge on the gene network rewiring upon *in vitro* reprograming with egg extracts remains elusive.

The question is still open in fish as to whether *Xenopus* egg extract can alter the course of somatic cells in culture, and if this treatment bears the potential to later on improve nuclear transfer in fish species. In a previous study, we demonstrated that goldfish fin cells in primary culture can incorporate *Xenopus* MII-egg extract molecules such as LaminB3 in their nucleus ^[31]^, thus providing evidence that egg factors can reach somatic cells chromatin. It has been described in mammals that the reprogramming effect of *xenopus* egg extracts requires a culture step of the cells, so that they can recover from the treatment and that the new cellular program can induce changes in gene expression ^[26]^. However, the treated cells in our former work were too fragile to be cultured, and the reprogramming consequences of the treatment had been impossible to study. In the present work, we have set up a procedure which allowed the survival and proliferation of the goldfish treated cells in culture. This enabled the analysis of their reprogramming extent. The use of nuclear transfer success as a mean to assess the extent of donor cell reprograming was excluded because of the multiparametric factors at stake in clone development success in fish. Indeed, embryonic failures are a combination of mitotic errors ^[32]^ and gene reprogramming defects ^[18,16]^ whose respective contribution is highly variable between clones. Therefore, the consequences are impossible to discriminate one from the other at the embryonic genome activation stage, when most clones fail to develop ^[11–13]^.

The aim of the present work was to investigate the response of goldfish somatic cells to treatment with *Xenopus* egg extract in culture, and to assess whether this treatment triggered some reprogramming events that would take place ahead of cell collection for nuclear transfer. Changes in gene expression were analyzed by an unbiased microarray approach, and specific networks associated with reprogramming were sought, in relation with the behavioral changes of the cultured cells. At the epigenetic level, changes in DNA methylation pattern of some candidate genes were also explored, namely the *pou2* and *nanog* genes that have differentially methylated promoters between fin cells and embryonic cells ^[33,34]^. Cells from primary fin culture were chosen over cell lines, because the former are closer to the original genetic background that is sought for regeneration of valuable fish genotypes by nuclear transfer.

## RESULTS

### Validation of fin cell exposure to *Xenopus* egg extract

The mesenchymal cell preparation and treatment that were set up in a previous study ^[31]^ included plasma membrane permeabilization with digitonin, permeabilized cell exposure to egg extract for 1h, and plasma membrane resealing (Supplementary Fig. S1). Penetration of egg factors per se was not tested here, because this would have required cell fixation. However, all cells displayed the phenotypic characteristics of permeabilized cells and egg extract-treated cells that were described previously ^[31]^: their nuclear membrane was more contrasted after permeabilization than in control cells, their adhesion capacity lessened during egg extract exposure and remained very low during the resealing step and the first 24 h of culture, and they all adopted a round and refracting morphology after resealing. Taken together, our observations indicate that all treated cell batches in the present study did incorporate egg extract.

### Treated cells need a suitable culture medium to survive and proliferate

The culture phase of the treated cells had to be mastered, so that the treated cells could undertake their new cellular program. When the conventional L15 medium was used, we observed that from the second day of culture on, many treated cells displayed a cubic shape (Fig 1, left picture) that contrasted with the much more elongated control cells (Fig 1, central picture). However, after 7 days of culture in L15, the treated cell density decreased, and many cells detached from the culture plate (Fig 1, left pictures), whereas control cells kept proliferating (Fig 1, inset d7). This inability of the treated cells to survive in L15 medium provided a first indication that the egg extract treatment had induced some changes in the treated cell physiology.

**Figure 1.**
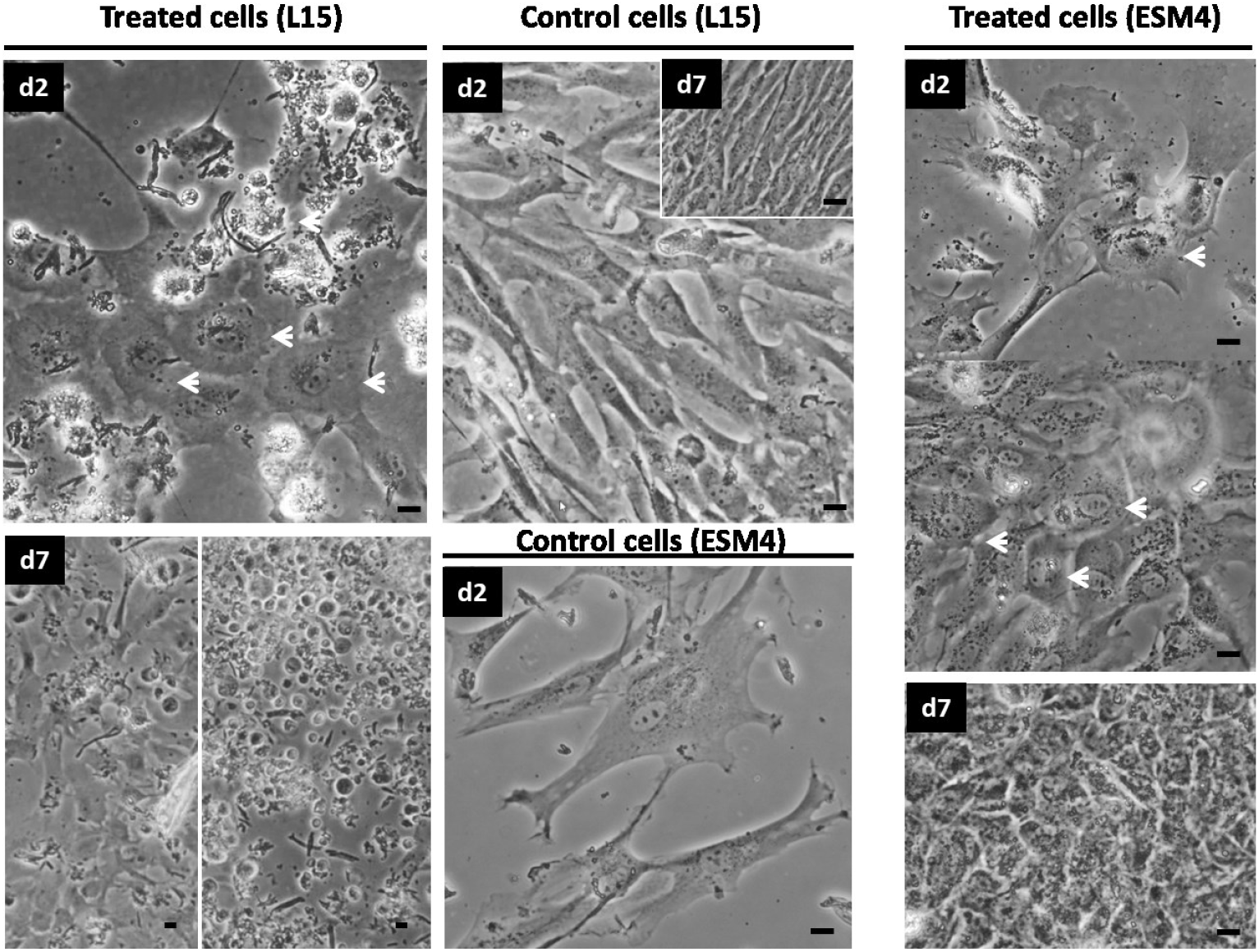
Characteristics of the cultured fin cells after Xenopus egg extract treatment. Treated cells were cultured in L15 (left) or ESM4 (right) media at 25 °C. Control cells (central) were neither permeabilized nor exposed to egg extract and they were cultured in the same conditions as the treated ones. Cell behavior and morphology were assessed by phase contrast microscopy after 2 (d2) and 7 (d7) days of culture. White arrows in d2 pictures: examples of cells with a round shape morphology, contrasting with the elongated control cells. Inset : typical morphology of confluent control cells after 7 days in L15 medium. At d7, treated cells quality was higher in ESM4 (d7, right) than in L15 (d7, left) medium. Pictures are representative of three experiments with different cells and egg extract batches. Scale bar = 10 μm.

Therefore, we sought for a culture medium that would sustain treated cells survival and proliferation, while maintaining their modified state. In mammals, somatic cells treated with egg extract were reported to be cultured in embryonic stem (ES) cells medium containing Leukemia Inhibitory Factor (LIF) and other complements aimed to prevent cell differentiation ^[29,25,27]^. However, fish ES-like cells are known to be independent from LIF (reviewed in ^[35]^). Furthermore, the maintenance of an undifferentiated state in zebrafish and medaka ES-like cultured cells was reported to require a medium enriched with fish serum and species-specific embryo extracts, known as ESM4 medium ^[35,36]^. We therefore tested the ESM4 medium enriched with goldfish embryo extracts (supplementary Table S1). After 2 days in this new culture medium, the cubic shape of the treated cells seen in L15 was maintained in ESM4 (Fig 1, right pictures). The elongated shape of the control cells was not changed either, indicating that ESM4 has no effect of its own on the shape of the cultured cells. Most interestingly, the treated cells cultured in ESM4 were able to proliferate over longer culture time compared to culture in L15 medium. They showed an increased cell density at day 7, and debris and floating cells were no longer observed (Fig 1, right picture). Additionally, they maintained their specific cubic morphology. This demonstrates further the favorable effect of ESM4 medium on the growth of the modified treated cells.

### Changes in gene expression eight days after egg extract treatment

#### Clustering of the differentially expressed genes (DEGs)

Analysis of the microarray data revealed that 2,286 goldfish genes out of the 52,362 genes on the microarray were differentially expressed between treated and control cells (fold change > 2). Additionally, hierarchical clustering analysis of the differentially expressed genes (DEGs) showed a clear segregation between treated and control samples (Fig 2, upper dendrogram). This demonstrated that the treated cells transcriptome was modified by egg extract treatment and that the consequences were detectable after 8 days of culture. Differentially expressed genes between treated and control cells showed a distribution into two clusters on the heatmap. Cluster I gathers 872 genes (encompassing 38 % of the DEG) that showed upregulation in the treated cultured cells. Cluster II comprises 1 414 genes (62 % of the DEG) that were down regulated in the treated cells. Genes in each cluster are listed in Supplementary Table S2.

**Figure 2.**
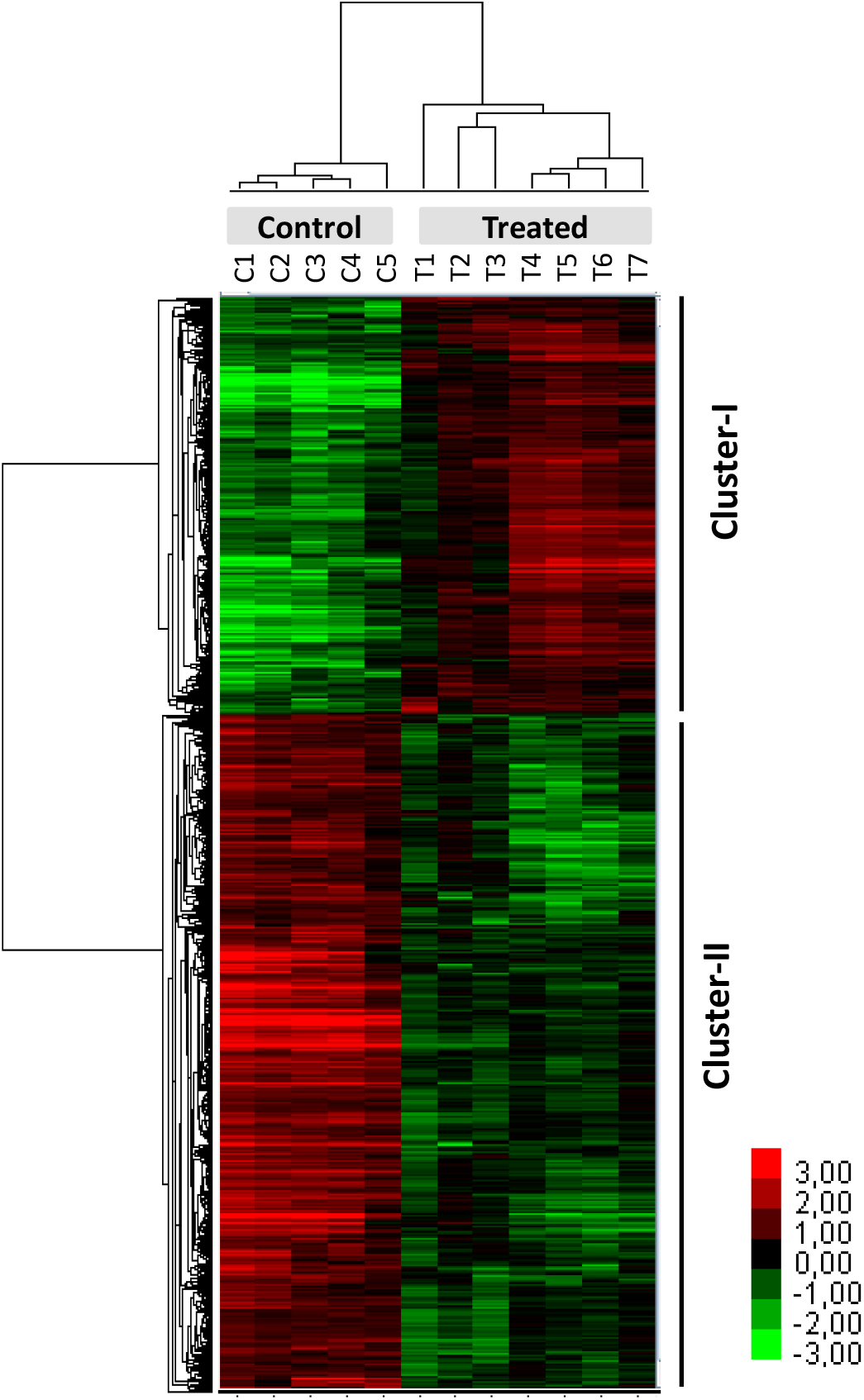
Hierarchical clustering analysis by unsupervised approach using 52,362 goldfish genes. Control: cultured control cells (C1-5); Treated: cultured treated cells (T1-7). Each row represents a single gene. Differentially expressed genes (Fold Change > 2; False Discovery Rate (FDR) < 0.05) between treated and control cells are shown on the heatmap (2,286 genes). Two clusters were identified. Cluster-I (872 genes) and cluster-II (1414 genes) contain the genes that were respectively up- and down-regulated in treated cells compared to control cells.

#### Segregation of the treated samples according to egg-extract batches

Although all treated samples segregated together and showed the same expression profile clustering, it is noteworthy that samples T1 to T3 segregated together, apart from the 4 other samples (T4-T7) (Fig 2, upper dendrogram). One possible explanation lies in the egg-extract batches that were used for the different samples. Indeed, the extracts were all prepared from freshly spawned MII stage eggs, each extract being obtained from the spawn of a different female. We cannot exclude that the individual extracts presented some quality variations one from another, notably because of the instability of MII stage in spawned eggs ^[37]^. In order to validate the egg extract stage, we used two MII markers: Greatwall whose phosphorylated forms prevent mitosis/meiosis exit ^[38]^, and Cyclin B whose degradation characterizes mitosis/meiosis exit. Egg extracts arrested at MII stage all displayed a specific western blot profile (supplementary Fig. S2): Greatwall (Gwl) was phosphorylated and stable over incubation time, and cyclin B (CycB) content was high and stable as well. Upon in vitro induction of MII exit by Ca^2+^, Greatwall was successfully dephosphorylated and cyclin B underwent degradation. Contrarily to these well-defined MII egg extracts, some egg extracts showed Greatwall dephosphorylation and Cyclin B degradation, indicating that they had initiated MII exit (MII late stage). Interestingly, the egg extracts used to treat T1 to T3 samples where in MII stage whereas those of T4 to T7 samples had initiated MII exit to some extent (Supplementary Fig. S2). As a conclusion, the sample segregation in the treated cells was likely related to the extract stages (MII and MII-late). This highlights the importance of a careful characterization of the *Xenopus* egg extracts. Although the sample number in each category was low, we still performed a fold change analysis between the two groups. We observed that 83% of the DEGs between MII and MII-late extract groups had low fold changes (< 6), and only 52 genes had fold changes above 6, among which only 9 genes were above 20. Besides, no significant or straightforward biological processes were identified via the GO terms analysis, and no marker gene of any specific biological significance emerged from a gene to gene scouting. To conclude, and within the limits of this small sampling, the egg extract stage did not thoroughly affect cellular response, and the two clusters of up- and downregulated genes were observed in all 7 treated cells batches irrespective of the egg extract that was used.

#### Gene Ontology (GO) analysis of the differentially expressed genes after egg extract treatment

GO analysis was a perquisite in order to process our DEG list into functions and biological significance. For this purpose, we had first to translate the goldfish gene identifiers into those of the closest species whose genome is well annotated in the GO databases, the zebrafish. This artificially reduced the number of DEG, because the zebrafish did not undergo the genome duplication reported in the *Cyprininae* sub-family to which goldfish belongs. Only 1533 zebrafish genes (i.e. 67% of total goldfish DEG) were retained for subsequent annotations in GO. Of these, 591 annotated genes were up-regulated (cluster I) and 942 annotated genes were down regulated (cluster II) in the treated cells. GO analysis conducted with the WebGestalt web tool ^[39]^ showed that biological regulation and metabolic process were the most represented terms (supplementary Table S3) among biological process GO terms. Surprisingly, no straightforward reprogramming processes such as chromatin remodeling, stem cells, transcription factors, or pluripotency could be emphasized in GO terms after a statistical over-representation analysis. However, several other biological process terms significantly enriched in the GO terms list deserve specific attention

#### Deregulation of TGFβ and Wnt signaling pathways after egg extract treatment

Our work is reporting gene expression variation, but GO databases and related publications on gene function report mainly protein functions. It is therefore the protein writing nomenclature that will be used in the following sections dedicated to GO interpretation. The most significant GO term obtained from the cluster of up-regulated genes is the cell surface receptor signaling pathway (Fig 3A). The data mapping showed that this GO term was linked to highly significant child GO terms that are transforming growth factor beta (TGFβ) receptor signaling pathway, and Wnt signaling pathway together with regulation of canonical Wnt signaling pathway. This result was consistent with the KEGG (Kyoto Encyclopedia of Genes and Genomes) analysis performed on the same set of DEG data, that also showed that both TGFβ and Wnt signaling pathways reached a significant level of enrichment among all the database terms (Fig 3B).To add on to the highlighting of these 2 specific pathways, we also observed an enrichment in the GO terms related to the MAPK / ERK cascade (Fig 3A), known to be one of the non-canonical pathways activated by TGFβ ^[40]^. Thus, the GO analysis based on the cluster of up-regulated genes clearly highlighted the TGFβ and Wnt signaling pathways as major ones being affected in the cells exposed to egg extract reprogramming factors.

**Figure 3.**
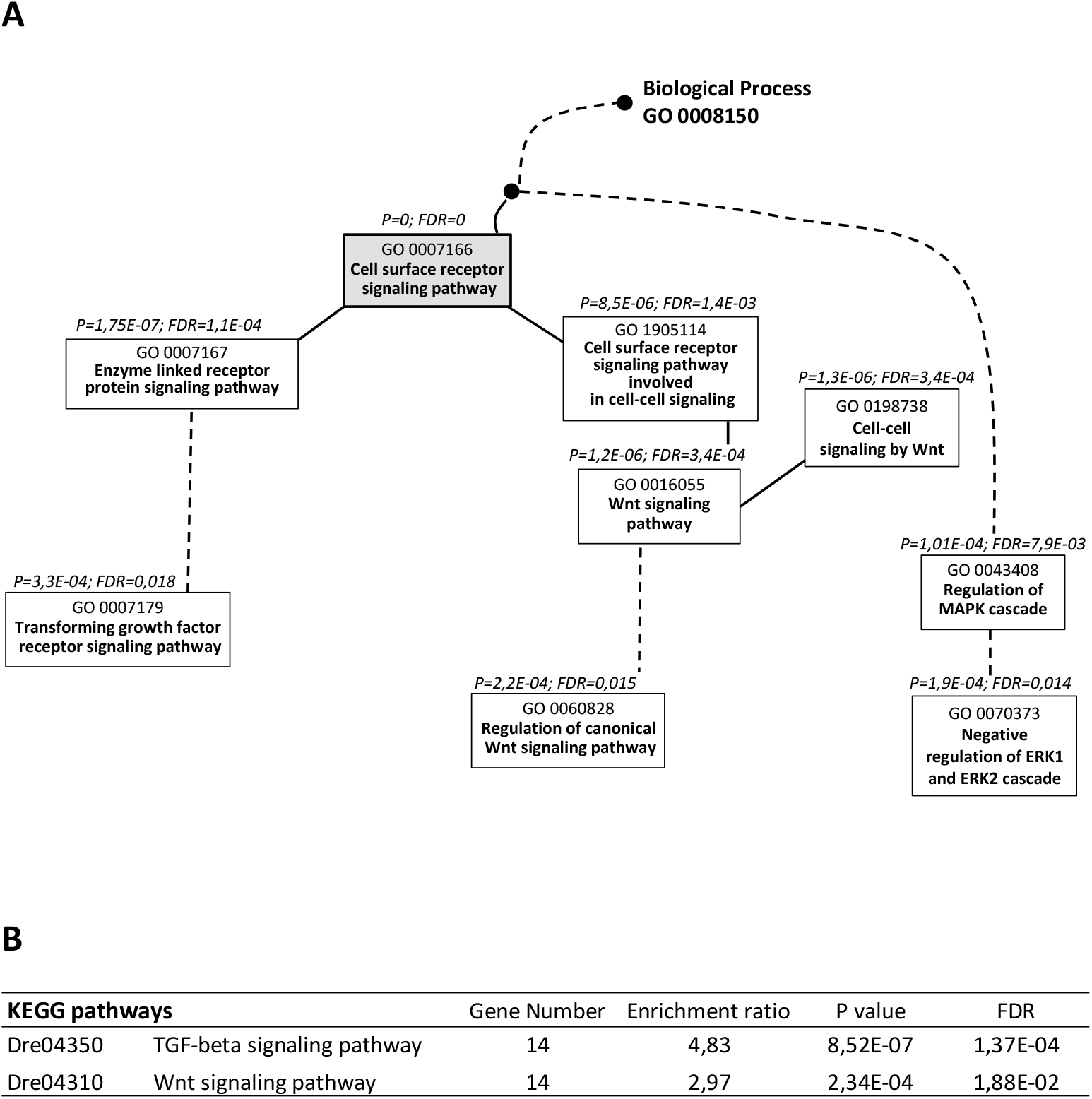
Gene Ontology (GO) flow diagram of the terms related to cell surface receptor signaling pathway (A) and KEGG pathways (B). The analysis was performed on the cluster of upregulated genes in treated cells (fold change >2) using WebGestalt web tool. The set of genes spotted on the microarray was used as the reference gene list. Black and dotted lines (in A) represent respectively direct and indirect connections between GO terms. Drexxxxx (in B) gives Danio rerio prefix of the KEGG identifier. For each GO term and KEGG pathway, p values (P) below 0.05 and false discovery rate (FDR) below 0.05 are indicated. Both A and B highlight the disturbance of the TGFβ and Wnt signaling pathways in response to egg extract treatment.

#### TGFβ signaling

TGFβ signaling is involved in numerous biological processes related to embryonic development. We then checked the actors of the TGFβ pathway present in our goldfish DEG list (irrespective of their up or down regulation). TGFβ belongs to the superfamily of the growth factors, divided into several subfamilies including TGFβs, and Bone Morphogenetic Proteins (BMPs). For TGFβ signal to be transduced, the TGFβ ligand binds type II receptors. Ligand - type II receptor complex triggers the recruitment of TGFβ type I receptor, and the dimerized receptors subsequently activates specific Smad proteins, able to induce transcription of the TGFβ target genes ^[41,42]^. Beyond signaling pathways involving Smads, known as canonical TGFβ pathways, other pathways independent of Smads are also controlled by TGFβ, including the MAPK Erk1 / ERk2 pathway identified above by GO analysis (Fig. 3A). We therefore analyzed the expression profile of these TGFβ actors and their biological partners.

We found that some TGFβ and BMP ligands together with type I receptors were upregulated in treated cells compared to controls (Table 1, TGFβ Effectors). However, this upregulation is unlikely stimulating the TGFβ signaling pathway, because the key actors binding TGFβ that are the TGFβ type II receptors did not change their expression pattern in treated cells. Besides, most other actors of the TGFβ signaling pathway identified in this study were affected in the direction of a TGFβ signaling inhibition in the treated cells, namely inhibitors upstream of TGFβ signaling that were upregulated in the treated cells (Table 1, TGFβ Inhibitors). Among them, we identified extracellular inhibitors (lft2, nog1, nog2, grem2a, grem2b) and membrane inhibitors (Bambia and Bambib) that are binding to TGFβ and BMP ligands. Such binding prevents TGFβ and BMP to attach to their own receptors, thereby preventing signal transduction activity ^[43,44]^. Beyond these inhibitors, we also found intracellular inhibitors (involved in TGFβ canonical signaling pathway) which included specific smads (smad6a, smad6b, smad7, smad9) and the ubiquitin ligase smurf2 (Table 1). The combined action of Smad7 and Smurf 2 is known to induce TGFβ type I receptor degradation by the proteasome ^[42,45]^, leading to inhibition of the TGFβ canonical pathway. Finally, spry1, sry4, and dusp6 genes, inhibiting the MAPK / ERK pathway (non-canonical TGFβ pathway), were also found upregulated in the treated cells (Table 1). In all, our gene to gene analysis of the expressional changes of TGFβ actors, including the non-canonical MAPK / ERK pathway, indicated that fin cells exposure to egg extract induced an overall inhibition of the TGFβ signaling pathway.

**Table 1.**
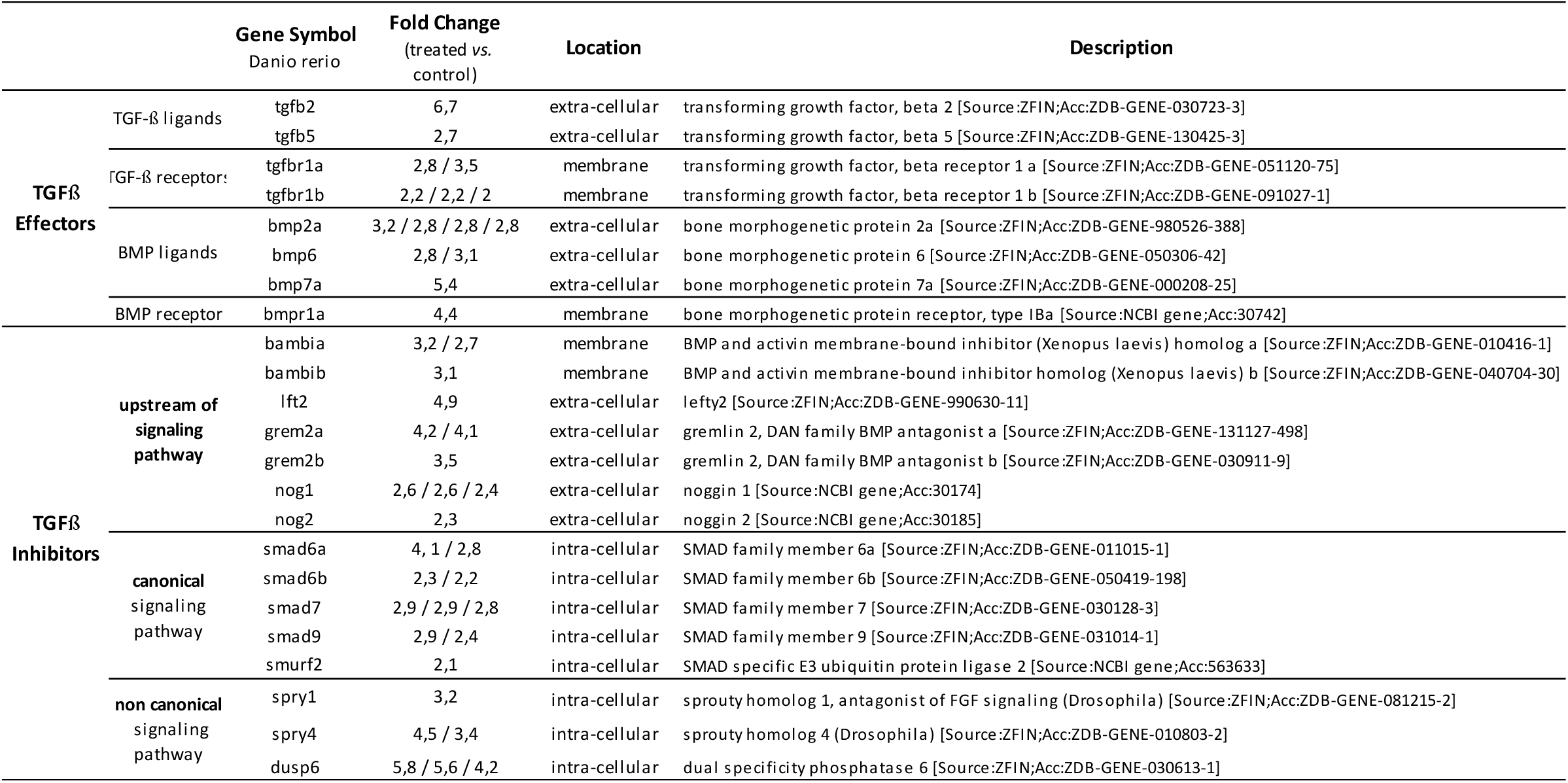
List of the TGFβ signaling actors that were upregulated in treated cells. For each Danio rerio gene symbol, Fold Change values are given for all the corresponding isoforms found in goldfish. Despite upregulation of some TGFβ effectors, upregulation of many inhibitors acting upstream and downstream of the signaling pathway are signing TGFβ inhibition in treated cells. Both canonical and non-canonical signaling pathways were affected.

#### Mesenchymal-epithelial transition

In mammals, one consequence of TGFβ signaling inhibition is the induction of a mesenchymal-epithelial transition (MET), considered to be a hallmark of iPSC early phase reprogramming, and described as crucial for reaching pluripotency ^[46–50]^. MET is characterized by the loss of mesenchymal markers and by the activation of genes determining epithelial fate ^[47]^. We therefore investigated whether inhibition of TGFβ signaling in our fish treated cells was also associated with changes in MET marker genes. We found that many mesenchymal marker genes were downregulated, among which several members of the collagen family, matrix-metallo protease (mmp9) and fibronectin (fn1) (Table 2). The fn1 gene was the most strongly affected (−44 fold change). This was associated with the concomitant upregulation of several epithelial marker genes such as cadherins (pcdh1 cadherin-like 1, pcdh12, cdh18, cdh24b), cytokeratins (krt15, krt18), and cell junction proteins such as pkp3b, cldn5a, tjp1a and cx43 (Table 2). Regarding the gap junction component cx43, it is known to be specifically enriched in epithelial cells and iPSCs, and its ectopic expression and gene upregulation has been associated with an increase in reprogramming efficiency by facilitating MET ^[51]^. Last, the transcription factor zeb1 known to induce EMT (epithelial-mesenchymal transition) ^[52]^, ie the reverse of the MET, was downregulated in treated cells (Table 2). In all, the observed expressional changes suggest the initiation of a MET program in the treated cells. This expressional profile is in accordance with the epithelial-like morphology reported above for the treated cells, which were more cubic than the elongated control cells. However, we observed from this DEG analysis and from qPCR analysis that one abundant mesenchymal marker, col1a1a, remained highly expressed in treated cells and was not differentially expressed between the two conditions (relative expression 125.0 ± 52.8 in treated cells, n=7; 123.1 ± 25.8 in control cells, n=8). This suggests that MET would be initiated but not terminated in our culture conditions.

**Table 2.**
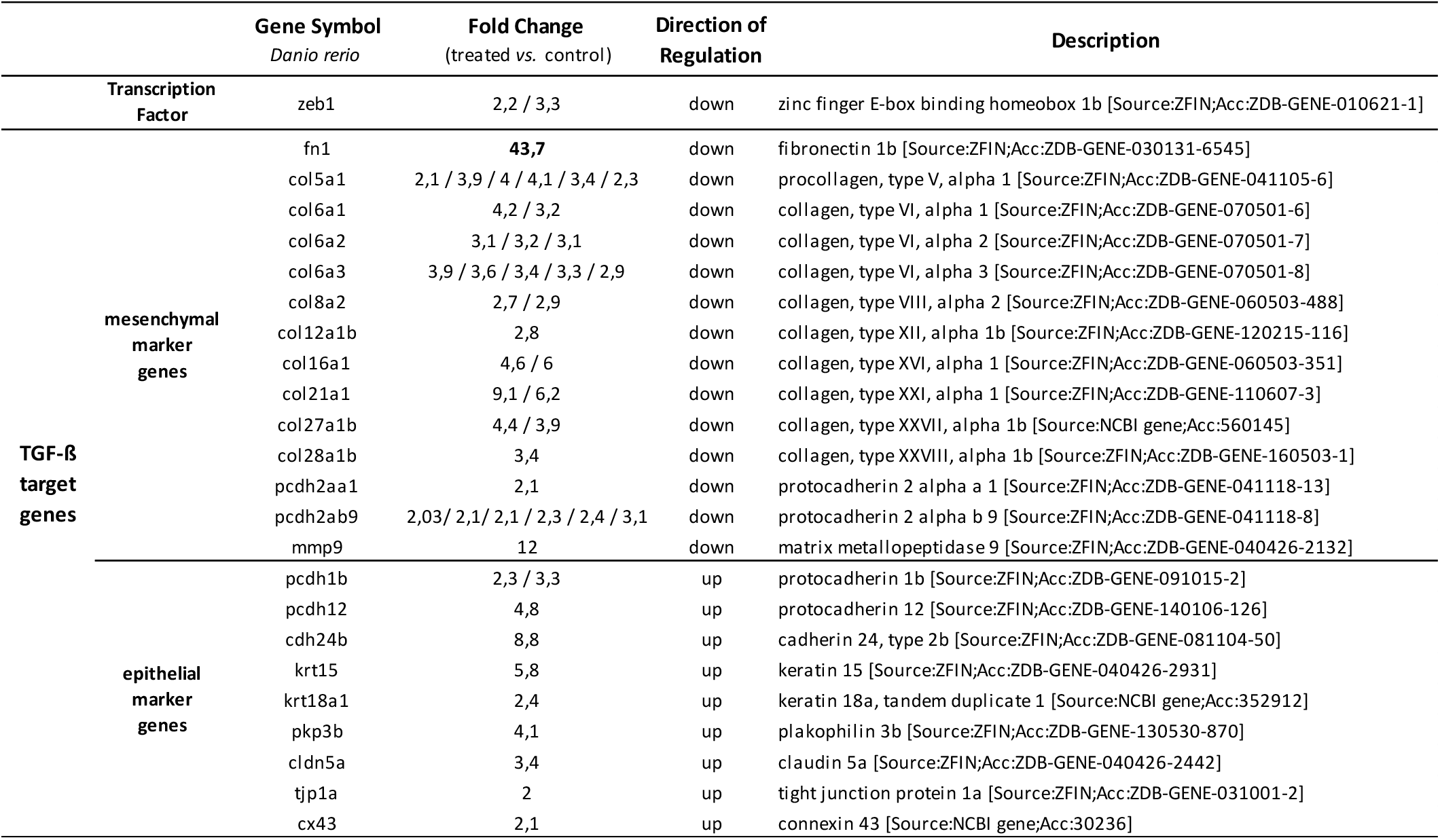
List of the TGFβ target genes related to mesenchymal-epithelial transition (MET) that were differentially expressed between treated and control cells. For each Danio rerio symbol gene, Fold Change values (up and down) are given for all the corresponding isoforms found in goldfish. Treated cells were characterized by the downregulation of several mesenchymal markers, especially fn1 and collagens, and by the upregulation of several epithelial markers, suggesting a MET.

#### Wnt signaling

The second signaling pathway whose terms were enriched in the GO analysis is the Wnt signaling pathway, and particularly the canonical one (Fig. 3). Because β-catenin is a key effector of Wnt signaling, the canonical pathway is referred to as Wnt/β-catenin signaling. The transduction of Wnt signal requires Wnt-induced activation of the receptors complex made of Frizzled (fzd) receptor and low-density lipoprotein co-receptor related 5 or 6 (LRP5/6). In other words, binding of the Wnt ligand to both receptors creates and activates the receptors complex. This initiates a series of molecular events that will protect cytosolic β-catenin from degradation. After nuclear import, β-catenin subsequently triggers the transcription of Wnt target genes by binding to transcription factors belonging to the T-cell factor/Lymphoid enhancer factor (Tcf/Lef) family ^[53,54]^.

Our gene to gene analysis of these Wnt-related actors revealed a strong deregulation of the Wnt/β-catenin signaling pathway in egg-extract treated cells (Table 3). Up-regulation of Wnt effectors combined with down- and up-regulation of inhibitors prevented the identification of a straightforward status for the Wnt signaling, be it an activated “on” or inhibited “off” status. In favor of an “on” status is the fact that some secreted Wnt ligands and fzd receptors were up-regulated in treated cells, the expression of the fzd10 receptor being especially strong. Moreover, extracellular Wnt agonists R-spondins (rspo2, rspo3), known to increase fzd receptors availability on the cell surface ^[53]^ and to stabilize the LRP5/6 co-receptors ^[54]^, were up regulated in treated cells. Additionally, down regulation of the extracellular inhibitors sfrp1a, sfrp2 and dkk1a ^[53]^ should be inducing a better availability of the Wnt ligand for fzd receptors.

**Table 3.**
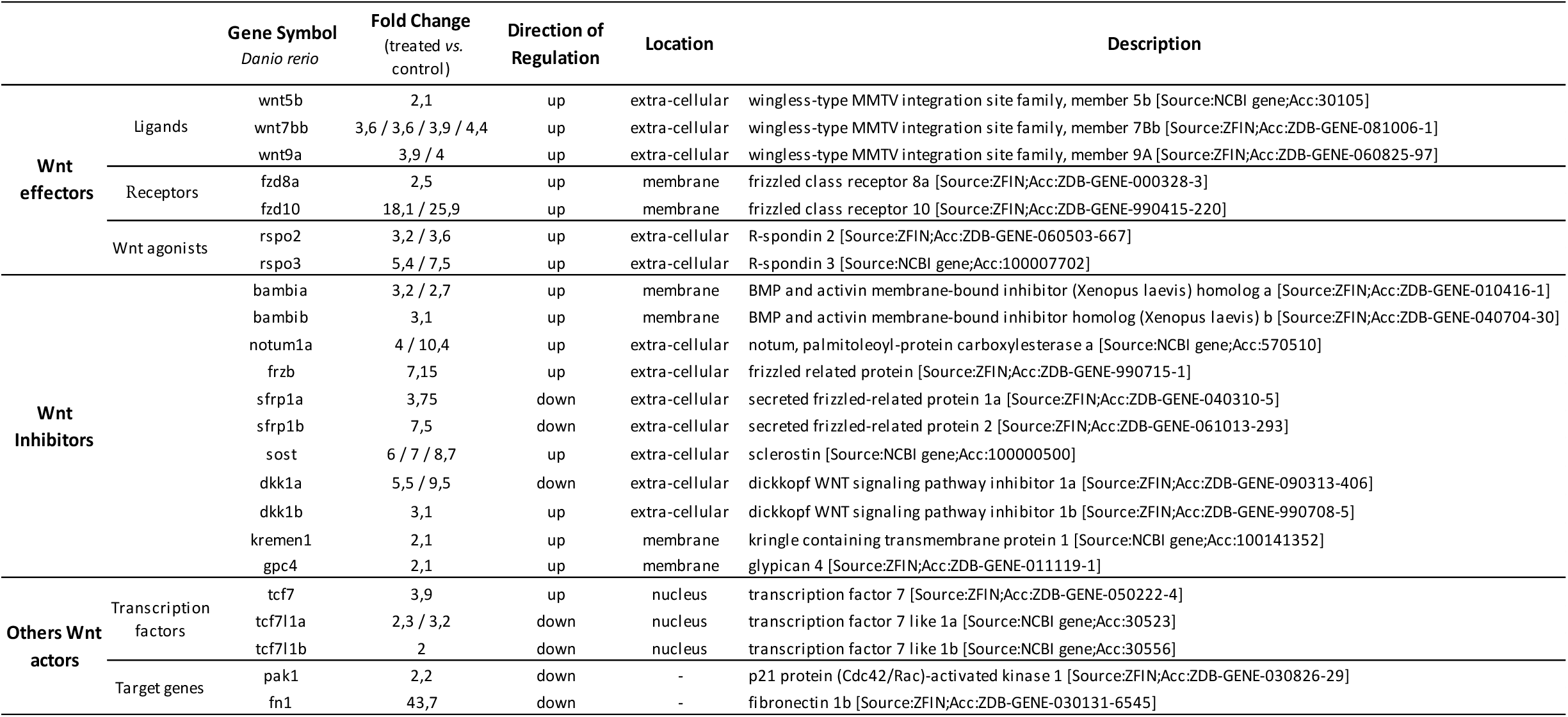
List of the Wnt/β-catenin signaling actors that were differentially expressed between treated and control cells. For each Danio rerio symbol gene, Fold Change values (up and down) are given for all the corresponding isoforms found in goldfish. The overall disturbance of the Wnt/ β-catenin signaling pathway in response to egg extract treatment, indicated by up/down regulation of effectors, inhibitors and transcription factors, tilts towards Wnt signaling inhibition.

However, other observations would be more in favor of an “off” status of the Wnt signaling. Firstly, the co-receptors LRP5/6 expression was not changed by the treatment. Despite increase in Wnt ligands and fzd receptors gene expression, LRP5/6 stability would stoichiometrically hamper the formation of the ternary proteic complex Wnt ligand / fzd receptor / LRP5/6 co-receptors that is essential for signal transduction. Secondly, many extracellular inhibitors upstream of the signaling pathway were upregulated in treated cells (Table 3). These included (i) notum1a and frzb, known to prevent Wnt ligand from binding to fzd receptor ^[54,55]^, (ii) sclerostin (sost) and dkk1b, which are blocking Wnt-fzd-LRP5/6 complex formation by interacting with LRP5/6 ^[53]^ and, (iii) kremen1, a membrane receptor which interacts with dkk1 to increase the removal of the LRP5/6 co-receptors from the cell surface by endocytosis ^[56]^. Finally, the last point concerns the Tcf/Lef transcription factors, known as Tcf1, Tcf3, Tcf4 and Lef1 in mammals, which control the Wnt signaling activity through transcription of the target genes. In this study, Tcf7 (orthologue of Tcf1 in mice) expression was upregulated in treated cells (Table 3). It is known that the action of Tcf1 is triggered by nuclear beta catenin levels (Grainger 2019). Tcf1 acts as a transcriptional activator of Wnt target genes in presence of beta catenin. Conversely, Tcf1 acts as a transcriptional repressor of Wnt signaling in absence of beta catenin (Grainger 2019). In our study, treated cells showed a considerable collapse of fn1 and to a lesser extent a downregulation of pak1, also known as p21 (Table 3). The downregulation of these Wnt target genes associated with Tcf7 upregulation suggest that Tcf7 would act rather as a repressor in treated cells due to low beta catenin levels. All these observations reinforce the hypothesis that the deregulation of the Wnt signaling in treated cells would be rather in “off” configuration.

#### Altered cell adhesion of the treated cells and link with the deregulation of TGFβ and Wnt signaling pathways

In addition to the change in treated cells morphology, we also reported above a change in their behavior in culture. The cells showed a highly reduced ability to adhere throughout the culture process, and this could be due to changes in some gene expression. And indeed, from the GO analysis of all DEGs between treated and control cells, one biological process GO term highlighted the cell adhesion process (GO: 0007155; *P value*=3.2560E-08; FDR=1.17E-05). Furthermore, fibronectin (fn1) is a major protein of the extracellular matrix and it provides highly adhesive capacity to the cells by interaction with integrin transmembrane receptors ^[57]^. As shown above, this actor was one of the most highly downregulated gene in our conditions, and this reduced expression is known to be deeply interconnected with the observed inhibition of the TGFβ and Wnt signaling pathways (Table 2, Table 3) ^[58]^. In all, the poor adhesion capacity of our treated cells is another indication of the undergoing reprogramming event triggered by changes in the TGFβ and Wnt signaling pathways expression.

### Some pluripotency markers remained silent in the treated cells

The process of somatic reprogramming in iPSCs is generally encompassing two phases ^[59]^: (i) an early or initiation phase during which the somatic cells undergo a MET, lose their mesenchymal characteristics and develop an epithelial phenotype and, (ii) a late maturation phase allowing the reactivation of the pluripotency network. In order to characterize further the reprogramming extent of the treated fin cells, we focused on some marker genes related to pluripotency, previously characterized in goldfish during early development: *pou2* (*pou5f3* in zebrafish, *oct4* in mammals), *nanog, sox2* and *c-myc* ^[33,34,60]^. We observed that none of these genes were identified among the DEGs, and their expression levels remained undetectable on the microarray. These observations were confirmed by qPCR that showed that *pou2, nanog, sox2* and *c-myc* expression was below detection threshold in both treated and control cells.

It was shown previously in goldfish that *nanog* and *pou2* silenced status in fin cells is associated with the hypermethylation of a CpGs locus in their promoter region ^[33,34]^. We also showed recently that after nuclear transfer with non-treated fin cells, these loci underwent a partial and stochastic demethylation in the developing clones ^[16]^. This indicated that embryonic reprogramming relaxed the DNA methylation status of these marker regions to some extent. The methylation profile of *nanog* and *pou2* promoter regions in our treated cells was therefore analyzed, to assess whether some DNA demethylation took place at these marker sites after *xenopus* egg treatment. This would indeed be a necessary step ahead of any transcription reenabling of these 2 genes. Analysis of the CpG sites in *pou2* and *nanog* promoter regions revealed that they did not underwent any significant demethylation in treated cells (Supplementary Fig. S3). Although the methylation of some CpG sites was lower in treated cells compared to controls, there was no significant differences in the overall DNA methylation rate of *pou2* and *nanog* promoter regions. This indicates that no significant remodeling of the DNA methylation of *pou2* and *nanog* pluripotency genes was triggered following egg-extract treatment, even if some minor variations were observed. In all, the silenced status of pluripotency marker genes associated with the absence of significant DNA methylation remodeling support the idea that treated cells would have been only partially reprogrammed by *Xenopus* egg-extract treatment. Our cells would not have reached the maturation phase of reprogramming characterized by Oct4 or Nanog and Sox2 re-expression as observed in mammalian somatic cells.

### Alteration of *de novo* lipid biosynthesis in response to egg-extract treatment

Regarding the cluster of downregulated genes, the GO biological processes the most significantly affected by egg-extract are related to lipid metabolism (Fig 4A). Child GO terms targeted biosynthesis of steroid including cholesterol, and biosynthesis of unsaturated fatty acid. This was consistent with KEGG analysis showing the enrichment of the biosynthesis pathways of steroids, unsaturated fatty acids as well as the pathway of fatty acid metabolism (Fig 4B). In this process, acetyl-CoA represents the main precursor for de novo lipid biosynthesis. Produced in the mitochondria after glycolysis, acetyl-coA has to be metabolized into citrate so that it can exit the mitochondria. Once in the cytoplasm, citrate is then converted into lipogenic acetyl-CoA (see the molecular actors of lipogenesis in ^[61]^). A detailed analysis of lipid metabolism genes showed a downregulation of several genes involved in the cytosolic synthesis of acetyl-CoA i.e. slc25a1b, a key mitochondrial transporter of citrate, aclya, which converts cytoplasmic citrate to acetyl-CoA, acss2, which produces acetyl-CoA from acetate, and the acyl transferase acat2 (Table 4). Lipid biosynthesis is also controlled by srebf1/2 transcription factors, whose expression was downregulated in our treated cells. The target genes of these transcription factors were downregulated as well. These included key enzymes for biosynthesis of cholesterol (hmgcs1, hmgcra1, msmo1, fdft1, cyp51, dhcr7) and fatty acid (fasn, sdc, elov1a, elov2, elov5, elov6) (Table 4). Overall, our results clearly indicate that the treated cells have strongly reduced their *de novo* lipid biosynthesis compared to control cells.

**Table 4.** List of the genes associated with lipid biosynthesis that were downregulated in treated cells. For each *Danio rerio* symbol gene, Fold change values are given for all the corresponding isoforms found in goldfish. *: transcription factors involved in the regulation of fatty acid and cholesterol *de novo* synthesis.

|  | **Gene**<br>**Symbol**<br>*Danio rerio* | **Fold Change**<br>(Treated vs. control) | **Description** |
| --- | --- | --- | --- |
| **Acetyl-CoA**<br>**synthesis** | slc25a1b | 6,6 / 11,7 / 14,2 | slc25a1 solute carrier family 25 member 1b [Source:NCBI gene;Acc:795332] |
|  | aclya | 4,4 / 4,5 | ATP citrate lyase a [Source:ZFIN;Acc:ZDB-GENE-031113-1] |
|  | acss2l | 6,63 / 32,6 | acyl-CoA synthetase short chain family member 2 like [Source:ZFIN;Acc:ZDB-GENE-130530-723] |
|  | acsl3a | 59,8 | acyl-CoA synthetase long chain family member 3a [Source:ZFIN;Acc:ZDB-GENE-050420-181] |
|  | acat2 | 3 / 5 | acetyl-CoA acetyltransferase 2 [Source:ZFIN;Acc:ZDB-GENE-990714-22] |
| **Fatty acid**<br>**biosynthesis** | fasn | 2,8 / 2,8 / 3,4 | fatty acid synthase [Source:ZFIN;Acc:ZDB-GENE-030131-7802] |
|  | scd | 6,5 / 6,9 / 7,4 | stearoyl-CoA desaturase (delta-9-desaturase) [Source:ZFIN;Acc:ZDB-GENE-031106-3] |
|  | fads2 | 32,2 / 34,8 / 35 / 35,6 | fatty acid desaturase 2 [Source:NCBI gene;Acc:140615] |
|  | elov1a | 2,2 / 2,4 | ELOVL fatty acid elongase 1a [Source:ZFIN;Acc:ZDB-GENE-041010-66] |
|  | elov2 | 3 / 22,3 | ELOVL fatty acid elongase 2 [Source:ZFIN;Acc:ZDB-GENE-060421-5612] |
|  | elov5 | 2,8 / 5,5 | ELOVL fatty acid elongase 5 [Source:ZFIN;Acc:ZDB-GENE-040407-2] |
|  | elov6 | 2,1 / 9,6 | ELOVL fatty acid elongase 6 [Source:NCBI gene;Acc:317738] |
| **Cholesterol**<br>**biosynthesis** | hmgcs1 | 13,2 / 12,4 | 3-hydroxy-3-methylglutaryl-CoA synthase 1 (soluble) [Source:ZFIN;Acc:ZDB-GENE-040426-1042] |
|  | hmgcra | 5,1 / 5,2 | 3-hydroxy-3-methylglutaryl-CoA reductase a [Source:ZFIN;Acc:ZDB-GENE-040401-2] |
|  | msmo1 | 22,6 | methylsterol monooxygenase 1 [Source:NCBI gene;Acc:406662] |
|  | fdft1 | 6,3 / 6,7 / 6,7 | farnesyl-diphosphate farnesyltransferase 1 [Source:ZFIN;Acc:ZDB-GENE-081104-242] |
|  | cyp51 | 10,6 / 10,9 | cytochrome P450, family 51 [Source:NCBI gene;Acc:414331] |
|  | dhcr7 | 8,4 / 8,8 | 7-dehydrocholesterol reductase [Source:NCBI gene;Acc:378446] |
|  | dhcr24 | 8,1 | 24-dehydrocholesterol reductase [Source:ZFIN;Acc:ZDB-GENE-041212-73] |
|  | cyp7a1 | 4,6 / 5,5 | cytochrome P450, family 7, subfamily A, polypeptide 1 [Source:ZFIN;Acc:ZDB-GENE-040426-1296] |
|  | mvda | 4,4 | mevalonate (diphospho) decarboxylase a [Source:NCBI gene;Acc:492781] |
| **Transcription**<br>**factors \*** | srebf1 | 3,3 / 4 | sterol regulatory element binding transcription factor 1 [Source:ZFIN;Acc:ZDB-GENE-090812-3] |
|  | Srebf2 | 2,6 / 2,8 | sterol regulatory element binding transcription factor 2 [Source:NCBI gene;Acc:100037309] |

**Figure 4.**
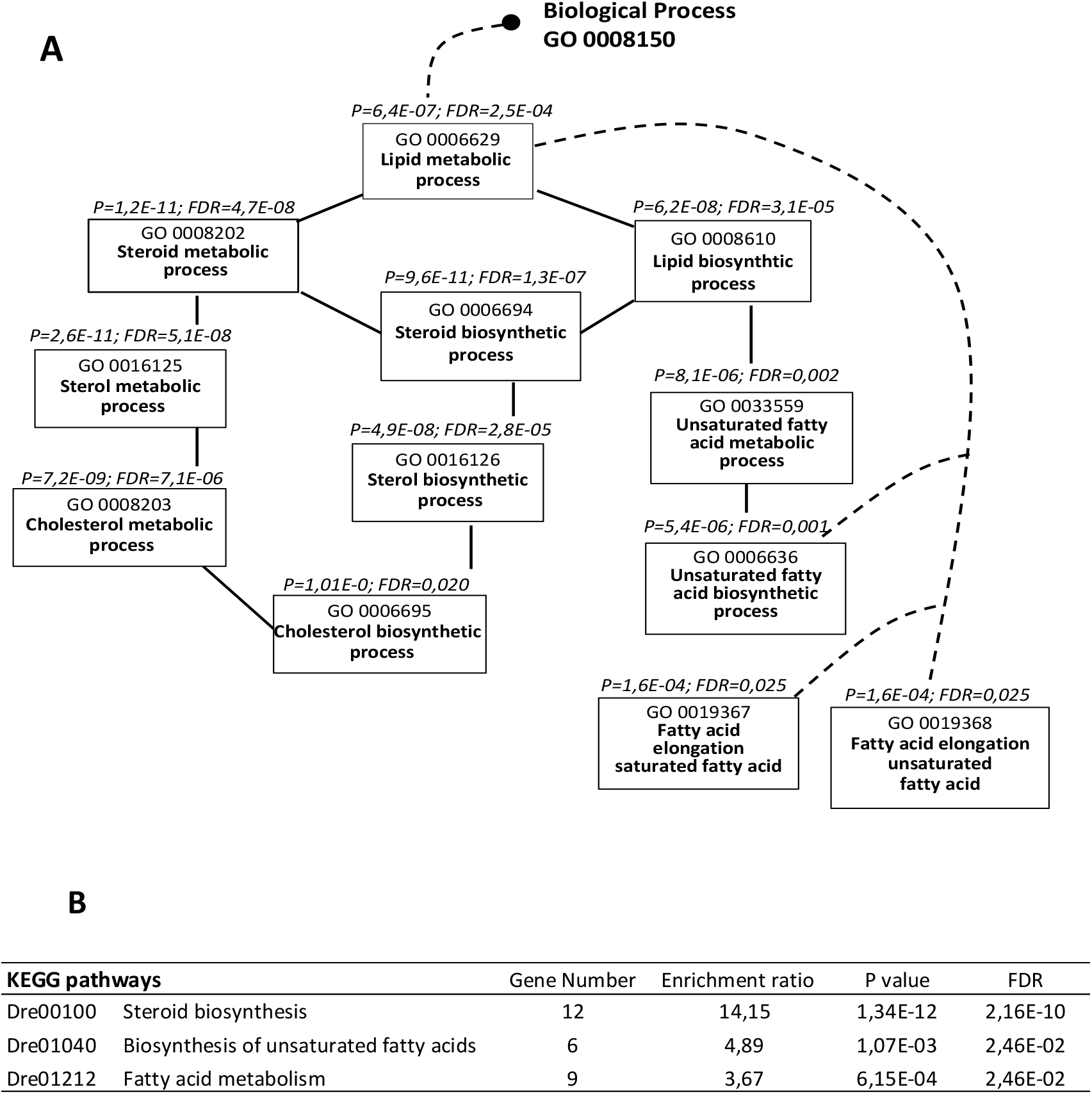
Gene Ontology (GO) flow diagram of the terms related to lipid metabolic process (A) and KEGG pathways (B). The analysis was performed on the cluster of genes downregulated in treated cells (fold change >2) using WebGestalt web tool. The set of genes spotted on the microarray was used as the reference gene list. A: The black and the dotted lines represent respectively direct and indirect connections between GO terms. B: Dre, Danio rerio prefix of the KEGG identifier. For each GO term and KEGG pathway, p values (P) below 0.05 and false discovery rate (FDR) below 0.05 are indicated. Both A and B highlight the disturbance of lipid metabolism after egg extract treatment, and specifically cholesterol and fatty acid biosynthesis.

## DISCUSSION

In this work, we explored to what extent fin somatic cells in culture could be modified by exposure to *Xenopus* egg-extracts. Our objective was to relax these differentiated cells so that they would be better fitted for reprogramming after nuclear transfer in fish. We showed that the treatment triggered some phenotypic changes of the cells, namely reduced adhesion capacity, adoption of a cubic shape morphology (epithelial-like), and reliance on the ESM4 medium for survival and proliferation. At the transcriptomic level, the thorough coverage of all goldfish transcripts and the number of biological replicates in our microarray analysis allowed us to identify two clusters of differentially expressed genes. We showed by GO analysis that cell surface receptor signaling pathways and lipid metabolism were the most significant terms that stood out from the list of these genes. Actors of the TGFβ and Wnt/β-catenin signaling pathways were altered in a pattern indicative of their inhibition, and this was combined with markers of a MET initiation. These changes were also associated with the lack of restoration of pluripotent markers activity and of their promoter demethylation, and with a reduction of lipid biosynthesis.

### Evidences of a reprogramming priming of the cells treated by *xenopus* egg extract

Reprogramming of somatic cells into a less differentiated state encompasses a series of molecular changes whose sequence includes the downregulation of somatic markers and of some signaling networks, induction of MET, and activation of early pluripotency markers ^[62,63]^. The TGFβ signaling inhibition observed in our treated cells, including that of the non-canonical MAPK / ERK pathway, would be one preliminary step in this reprogramming process. Indeed, experimental inhibition of TGFβ signaling was shown to cooperate in the reprogramming of murine fibroblasts into iPSCs ^[46,49,64]^. Furthermore, ERK inhibition was also shown to be an early molecular signature of somatic cell reprogramming in this model species ^[65]^, and inhibition of both TGFbeta receptors and ERK ^[48]^ also improved fibroblast reprogramming ^[66]^. Beyond these examples, our treated cells also displayed numerous MET markers, including the decreased expression of extracellular matrix genes, such decrease being known to mediate fibroblast reprogramming ^[67]^. Taken as a whole, our results lead us to propose that the egg extract would have initiated a reprogramming of the fin cells by directing them towards a MET via inhibition of TGFbeta signaling.

It is known that Wnt signaling deregulation is also triggering cellular reprogramming. In mice, it was shown that Wnt off state matches the early phase of iPSCs reprogramming of embryonic fibroblast ^[68]^. The off state of this signaling pathway observed in our study would indicate that our cells are in an early stage of reprogramming. One notable consequence of Wnt off state is that among the downstream target of the Wnt pathway, pak1 (orthologous to mammalian P21) was downregulated. This senescence actor was shown to be a barrier to iPSCs reprograming ^[69,70]^ and to MET. We therefore infer that its inhibition was favorable to MET and reprogramming in our treated cells.

### Incomplete reprogramming of the treated cells

As explained above, the treatment applied to our culture cells was intended to increase somatic cell plasticity towards reprogramming after nuclear transfer. Thus, our treatment with X*enopus* egg extracts remained within physiological limits, and it could not be expected to be as thorough as after reprogramming into iPSCs. Several indicators in our study showed that indeed, the treated fin cells were not entirely changed in their transcriptomic profile. Namely, although several collagens underwent a reduced expression in the treated cells, col1a1a abundantly expressed in fin cells ^[71]^ remained highly expressed after cell treatment. Furthermore, we failed to detect any re-expression of the canonical pluripotency markers that are *pou2, sox2, nanog* and *c-myc*. This is at odd with the Oct4 re-expression induced with a similar treatment in porcine or human cultured cells ^[26,28]^. Because of the stochastic re-expression of these genes described by these authors and reviewed in ^[72]^, we infer that we did not asses these markers expression in the right reprogramming window, or that our cells may still be in the initiation or intermediate phase of reprogramming. This emphasize the importance of a transcriptomic approach for a comprehensive assessment of cell reprogramming status.

### Reduced lipid metabolism in the treated cells

We showed that the whole cluster of downregulated genes induced the high significance of GO terms related to lipid metabolism, and numerous actors of the lipid biosynthesis were downregulated after *Xenopus* egg treatment. This questions the role of lipids in our cellular reprogramming scheme. Indeed, studies on iPSCs indicate on the contrary that increased lipid biosynthesis is favorable to MET and reprogramming ^[73,74]^, and that conversely, inhibition of fatty acid biosynthesis blocks mouse embryonic fibroblast reprogramming to iPSCs ^[75]^. It was shown that large amounts of lipids are consumed during the reprogramming process, as judged by the decreasing number of lipid droplets per cell between the early and late stages of reprogramming ^[75]^. This would indicate that lipid biosynthesis upregulation is intended to provide for additional energetic resources to the cells undergoing reprogramming. This hypothesis was explored in porcine iPSCs ^[76]^, and it was demonstrated that supplementation of the culture medium with triglycerides, free fatty-acids, phospholipids and cholesterol improved the reprogramming of embryonic fibroblasts by promoting MET.

Although we unambiguously showed in our study that egg-extract treatment failed to remodel the lipid metabolism of fin cells according to iPSC pattern, we cannot exclude that lipid biosynthesis downregulation was a response of the treated cells to the enriched ESM4 culture medium. In the conventional L15 medium, the treated cells died after a few days. It means that if the treated cells suffered endogenous lipid exhaustion, the lipids provided by their short-term exposure to *Xenopus* egg extracts were not able to compensate for such losses. On the contrary, subsequent culture in the ESM4 medium containing extracts from goldfish embryos at 55 hours post fertilization stage may have provided for the required energetic substrates. Fish embryos at this stage are indeed highly enriched in cholesterol, phosphatidyl choline and triglycerides ^[77]^. This means that ESM4 may have provided the same MET-favorable environment as in ^[76]^. In all, we may infer that the exogenous lipid supply via ESM4 would have met the need of the treated cells, possibly supporting the MET requirements, and that as a response, the lipid anabolism of the cells was reduced, leading to the observed downregulation of the corresponding genes.

To conclude, the treatment of fish fin cells with *Xenopus* egg extract and subsequent culture in ESM4 induced phenotypical and expressional changes signing the induction of a reprogramming process via MET. The identified reprogramming markers encompassed the TGFβ and Wnt/β-catenin signaling pathways inhibition, the cubic cell shape, and the lessened cell adhesion capacity. We also provided evidences that the reprogramming was obviously incomplete, as attested by the lack of pluripotency markers re-expression, and maintenance of one abundant mesenchymal marker. Taken as a whole, it appears that the fish somatic cells would be in early reprogramming phases, and that the treatment helped to release some of the reprogramming barriers present in our differentiated fin cells. This reprogramming priming by the *Xenopus* egg extract treatment could be a first favorable step towards expression reprogramming and chromatin remodeling after nuclear transfer.

## METHODS

### Goldfish fin cell preparation

Two-year-old goldfish (*Carassius auratus*), 60 g mean weight, were obtained from outdoor ponds at INRAE U3E experimental facility (Rennes, France), and maintained in recycled water at 14°C for several weeks. Caudal fins were collected on euthanized fish following the French animal welfare guidelines and under the French registration authorization n° 78-25 (N. Chênais). Fin cells were isolated and cultured according to ^[31]^. Briefly, fins were minced and digested with 2 mg/mL collagenase. Released fin cells were plated in supplemented L15 culture medium. After 24 hours, adhering epithelial cells were discarded while the supernatant enriched with slow adhering mesenchymal cells was collected. These cells have previously been shown to be the most suitable for nuclear transfer ^[71,78]^. After filtration and washing, the mesenchymal cells were seeded at 0.2 × 10^6^ cells per well in 24 well plates and cultured in L15 medium for 2 days (about 80 % confluence) until *Xenopus* egg treatment.

### *Xenopus* egg extract preparation and characterization

Unfertilized eggs from *Xenopus laevis* 2 years old females were obtained after hCG stimulation at the CRB Xenope facility (University of Rennes 1, agreement number: 35–238-42). Egg extracts were prepared as described previously ^[31]^. Laid eggs were crushed at 10 600 g for 20 min at 4°C and the extract was clarified at 10 600 g for 20 min at 4°C. The supernatant was collected, snap frozen in liquid nitrogen and stored at −80°C. A total of 7 individual spawns were collected, providing 7 batches of independent egg extracts with a protein concentration of 40 to 50 mg/mL and an osmolality of about 400 mOsm / kg.

Egg extract stage was characterized by western blot analysis using the mitotic markers Greatwall and Cyclin B as described in ^[31]^. For each egg extract, one fraction was immediately denatured in Laemmli buffer at 95 °C (3 min). A second fraction was incubated at 25°C for up to 2 h prior to denaturation, to mirror the time during which cells were treated with the egg extract. A last fraction was incubated with 0.8 mM Ca^2+^ at 25°C for up to 2 h to test its responsiveness to calcium-induced activation, before it was denatured. MII status of the 7 egg extracts was determined using rabbit polyclonal *Xenopus* anti-Greatwall and anti-Cyclin B (1:1000 each) according to ^[31]^.

### Somatic cell treatment and culture

Adherent mesenchymal cells were permeabilized with digitonin (30 ug/mL 2 min 4°C) before exposure to egg extract (1 h, 25 °C), according to ^[31]^. Cells were then incubated for 2 h in growth L15 medium supplemented with 2 mM CaCl2 (25 °C) to reseal the plasma membranes and then cultured at 25°C in ESM4 medium ^[35]^ (Supplementary Table S1). Culture medium was changed every 3 days. After 8 days, cultured cells were collected after trypsinization and snap-frozen in liquid nitrogen. Non-permeabilized cells were grown in L15 medium and used as controls. They followed the steps as the treated cells and were snap-frozen after 8 days.

### Microarray analysis

#### Microarray preparation and hybridization

Agilent 8×32K high-density oligonucleotide microarray (GEO platform no. GPL32340) was spotted with a set of 52,362 distinct goldfish oligonucleotides. Available goldfish NCBI sequences were blasted on the zebrafish genome, generating a list of zebrafish proteins identified as ENSDARP in the Ensembl database. The official symbol of each gene, its description and its Ensembl ID, called ENSDARG, were then extracted from the ENSDARPs using the Ensembl Biomart programm.

Total DNA and RNA of the cultured cells were extracted simultaneously after cell lysis in RNAsin (1 μL) in Tri-Reagent, according the instructions for Miniprep DNA/RNA Direct-zol column extraction kit (Zymo Research, R2081). RNA labeling and hybridization were performed according to the manufacturer’s instructions (Agilent “One-Color Microarray-Based Gene Expression Analysis (Low Input Quick Amp labeling)”). For each sample, 150 ng total RNA was amplified and labeled using Cy3-CTP. Yield (>825 ng cRNA) and specific activity (>6 pmol of Cy3 per μg of cRNA) of the obtained Cy3-cRNA were checked on Nanodrop. Cy3-cRNA (600 ng) from each sample was fragmented, and samples were hybridized on randomly chosen sub-arrays for 17 h at 65 °C. After microarray scanning (Agilent DNA Microarray Scanner, Agilent Technologies, Massy, France), data were obtained with the Agilent Feature Extraction software (10.7.3.1) according to the appropriate GE protocol (GE1_107_Sep09) and imported into GeneSpring GX software (Agilent Technologies, Santa Clara, CA, USA) for analysis. Data were published at the NCBI’s Gene Expression Omnibus ^[79]^ and are accessible through GEO series accession number GSE205854. Of the 16 cell samples laid on the microarray, only 12 samples passed the quality controls and were selected for analysis (n=5 control; n=7 treated with egg extract).

#### Differentially expressed genes identification and Gene Ontology analysis

The raw gene expression data were normalized and transformed into Log2 values using GeneSpring software (Agilent). Only genes displaying an expression value significantly higher than that of the background in at least 75% of the samples and in at least one of the two conditions were retained. Selection of differentially expressed genes relied on a Student’s t-test with false discovery rate (FDR) correction and a fold change > 2 was applied. The significance level was set to FDR < 0.05 and p-value < 0.05. The DEGs were then classified according to their expression profile by unsupervised hierarchical clustering using Cluster3 software and were visualized by TreeView software.

A gene ontology analysis was carried out on the DEGs of each cluster using WebGestalt web tool (AnaLysis web-based GEne SeT AnaLysis toolkit). In order to highlight the GO terms related to biological process that were significantly enriched, an over-representation analysis (ORA) was carried out on the gene IDs (ENSDARG) of each cluster. For this, each gene list of interest was compared to a background gene list corresponding to all the genes spotted on the microarray. ORA was also carried out to search for KEGG (Kyoto Encyclopedia of Genes and Genomes) pathways. The significance level was set to below an FDR 5 % and a p value of 0.05.

#### RTqPCR analysis

Confirmation of the expression pattern of several marker genes, namely *col1a1a, nanog, pou2, sox2, c-myca1* and *c-myca2*, were achieved by RTqPCR analysis according to ^[71]^. The Ct values were normalized using the endogenous *18S rRNA* control and the target mRNA relative abundance was calculated according the formula: 2^−ΔCt^ with ΔCt = mean Ct (*target gene*) – mean Ct (*18S rRNA*).

### DNA methylation analysis

Total extracted DNA was purified using the Genomic DNA Purification and Concentration Kit (Zymo Research, D4010) and quantified using the QubitTM dsDNA HS Assay Kit (Q32851, Invitrogen). DNA was treated with bisulfite using the EZ DNA Methylation-Gold kit (Zymo Research, D5006) and regions of interest were amplified according to ^[16]^. Methylation status of the targeted CpGs was calculated after pyrosequencing with PyroMark Q24 ID 2.5 software (QIAGEN). Bisulfite conversion of control cytosines was above 98 %.

## ACKNOWLEDGEMENTS

The authors thank T. Lorca and N. Morin (UMR 5237 CNRS, France) for helpful exchanges and for providing anti-*Xenopus* Greatwall and CyclinB antibodies. We thank J. Montfort (INRA LPGP) for providing zebrafish protein ID (ENSDARP) corresponding to goldfish NCBI sequences. Thanks also to Bernard Joseph from INRAE U3E and Pierre Lô Sudan from ISC INRAE LPGP, who took care of goldfish rearing. Hélène Jammes and Antoine Peigné performed the DNA methylation assay and analysis. This work benefited from the financial support of the PIA CRB Anim ANR-11-INBS-0003

## AUTHOR CONTRIBUTIONS STATEMENT

NC designed the study, developed the experiments, analyzed the data, co-wrote the manuscript and prepared the figures. ALC designed goldfish DNA microarray, conducted the microarray experiment and provided preliminary statistical analysis of the microarray data. BG provided the indispensable *Xenopus* eggs and contributed to the discussion. JJL provided expertise and help for the gene ontology analysis and discussion. CL conceived the study and co-wrote the manuscript. All authors read and approved the manuscript.

## DATA AVAILABILITY STATEMENT

Microarray data were published at the NCBI’s Gene Expression Omnibus (^[79]^ and are accessible through GEO series accession number GSE205854. Most other data obtained in this work were provided in the supplementary file. Any missing data or supplementary information should be asked to the corresponding authors who will answer the requests in a timely manner.

