## Supplementary information for "Reprogramming of fish somatic cells for nuclear transfer is primed by *Xenopus* egg extract"

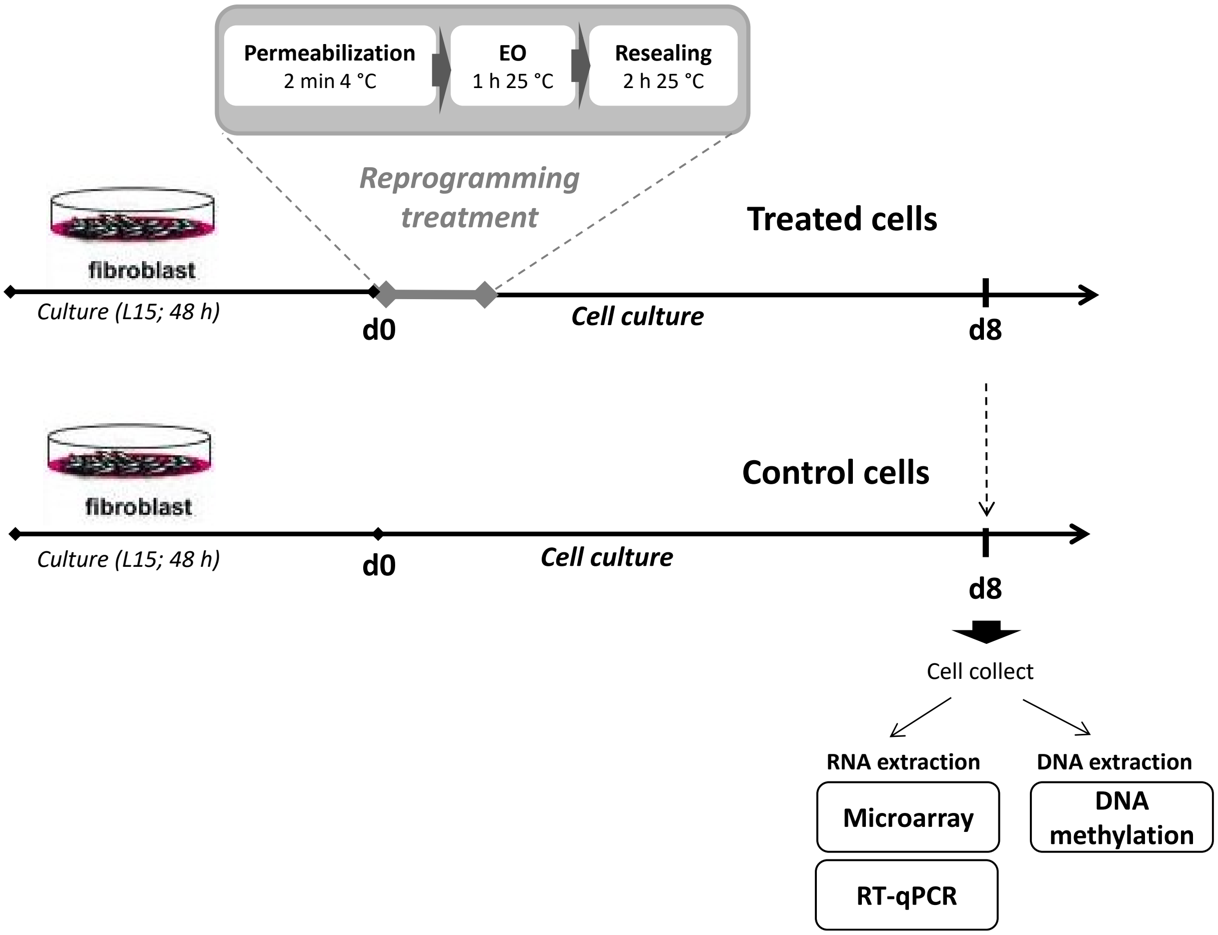


**Supplementary Fig. S1:** Diagram of the experimental approach for fish somatic cells reprogramming using *Xenopus laevis* egg extracts. Mesenchymal cells derived from fin primary culture were cultured 2 days in L15 medium before reprogramming treatment. This treatment included 3 steps i) plasma membrane permeabilization using 30 µg/ mL Digitonin, ii) exposure of the permeabilized cells to egg extracs and, iii) plasma membrane resealing. Treated cells were then cultured up to 8 days before RNA and DNA extraction for microarray, qRT-PCR experiments and DNA methylation analysis on candidate genes. Non-treated cells from the same batches but cultured in L15 medium were included as controls.

| Medium components | Content in ESM4 |
|---|---|
| L-Glutamine | 2 mM |
| Amphotericin B | 2.5 µg/mL |
| Gentamycin | 100 µg/mL |
| Pyruvic acid | 1 % |
| Non essential amino acid | 1 % |
| Sodium selenite | 2 nM |
| 2-mercaptoethanol | 100 µM |
| Fetal bovine serum | 10 % |
| Fish serum | 1 % |
| Human recombinant basic FGF | 8 ng/mL |
| Goldfish embryo extract | 1 embryo/mL |

**Supplementary Table S1:** Composition of ESM4 growth medium adapted to the culture of goldfish fin cells treated with *Xenopus* egg extract. The components listed in the table were added to the basic medium L15 medium supplemented with glucose (4.5 g/L), hepes (5 mM), sodium bicarbonate (2 mM) and adjusted to pH 7.3.

**Supplementary Table S2:** List of the differentially expressed genes between egg extract-treated and control cells.

Supl. Table S2_cluster-I genes_interestingID_mappingTable_wg_result1612179183.txt

Supl. Table S2 _cluster-II genes _ interestingID_mappingTable_wg_result1612178927.txt

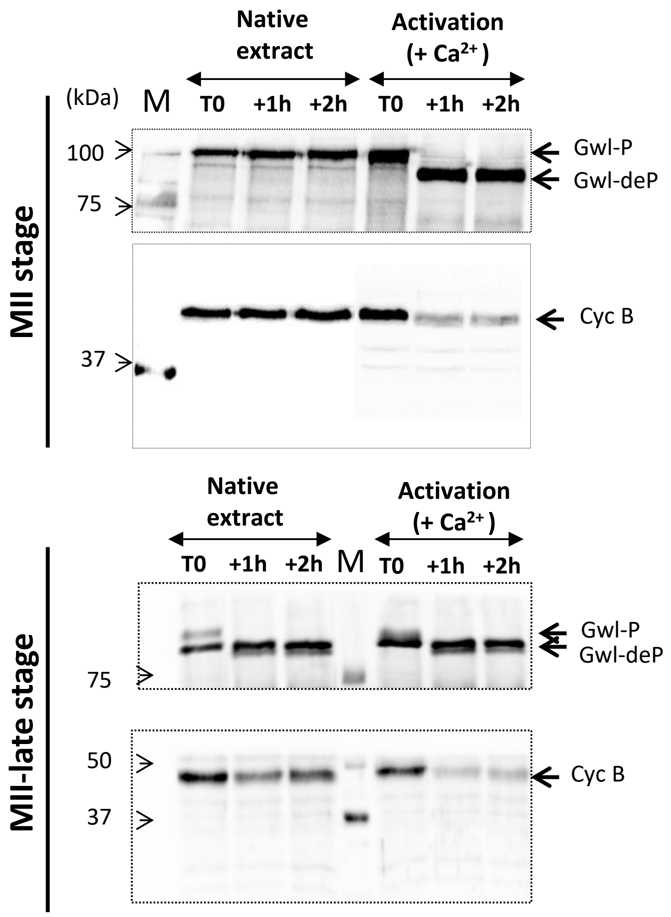


**Supplementary Fig. S2:** Mitotic status of egg extracts from *Xenopus* individual spawns: molecular characterization of their stage (MII or MII-late) by Western blot using Greatwall (Gwl) and Cyclin B (Cyc B) mitotic markers.

Endogenous Gwl-phosphorylation and Cyc B maintenance were monitored for each extract at different times: T0 (immediately after extract preparation), T+1h (corresponding to the 1 h exposure time of egg extract with cells) and T+2h to validate the stability of extract of MII-stage 5native extract). A control of mitotic exit corresponding to MII-late stage was also included for each extract after activation with $Ca^{2+}$ 0.8mM up to 2 h (Activation (+ $Ca^{2+}$)).

Note the existence of a strong phosphorylation differential of the Gwl marker between MII and MII-late stage extracts. MII stage extracts were characterized by a stable phosphorylation of Gwl (Gwl-P) for up to 2 h, whereas MII-late stage extracts displayed dephosphorylated Gwl (Gwl-deP). Besides, Cyc B was stable in the MII stage extracts for up to 2h and degraded in the MII-late stage extracts from the 1 h incubation on.

The profile of MII and MII-late stage egg extracts showed here are representative of the extracts used to treat cells (T1 - T3 samples and T4 – T7, respectively).

**Up-regulated genes in treated cells**

| GO identification | GO Terms | Gene number |
|---|---|---|
| GO:0065007 | biological regulation | 166 |
| GO:0008152 | metabolic process | 125 |
| GO:0050896 | response to stimulus | 118 |
| GO:0032501 | multicellular organismal process | 114 |
| GO:0032502 | developmental process | 110 |
| GO:0007154 | cell communication | 101 |
| GO:0051179 | localization | 67 |
| GO:0016043 | cellular component organization | 55 |
| GO:0008283 | cell proliferation | 17 |
| GO:0040007 | growth | 15 |
| GO:0051704 | multi-organism process | 9 |
| GO:0000003 | reproduction | 6 |

**Down-regulated genes in treated cells**

| GO identification | GO Terms | Gene number |
|---|---|---|
| GO:0065007 | biological regulation | 213 |
| GO:0008152 | metabolic process | 191 |
| GO:0050896 | response to stimulus | 152 |
| GO:0032502 | developmental process | 145 |
| GO:0032501 | multicellular organismal process | 144 |
| GO:0007154 | cell communication | 126 |
| GO:0051179 | localization | 82 |
| GO:0016043 | cellular component organization | 56 |
| GO:0008283 | cell proliferation | 16 |
| GO:0051704 | multi-organism process | 12 |
| GO:0040007 | growth | 11 |
| GO:0000003 | reproduction | 1 |

**Supplementary Table S3:** Gene ontology of differentially expressed genes between egg extract-treated and control cells. Distribution of biological process GO terms related to up-regulated and down-regulated genes.

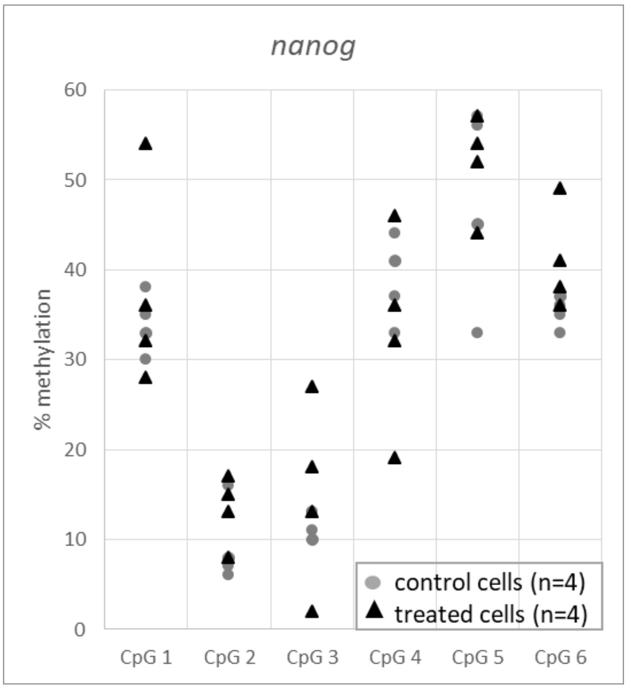


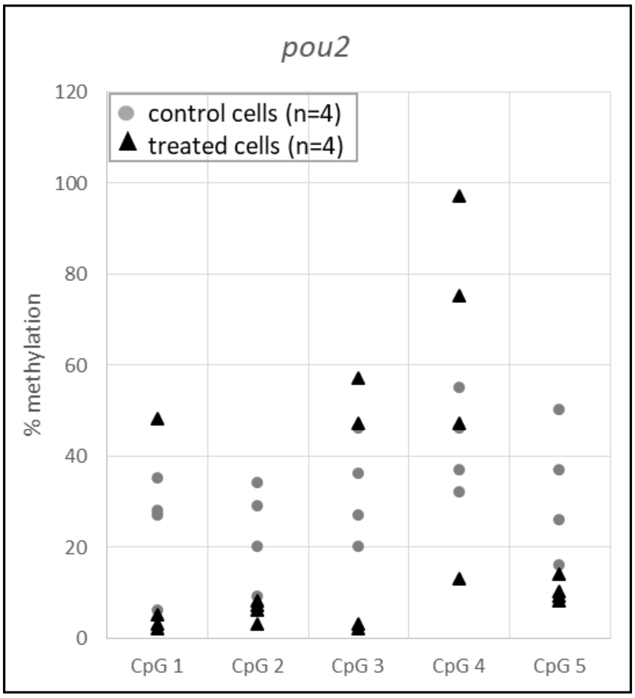


**Supplementary Fig. S3:** DNA methylation status of several CpG sites in the promoter regions of nanog (upper figure) and pou2 (lower figure) genes in relation to cell treatment. For nanog, CpG1, 2, 3, 4, 5, 6 are located respectively at the sites -324, -270, -217, -164, -134, -122 upstream of the putative transcription start site. Their average methylation status was 27.4 ± 3.3 % (control) and 32.0 ± 2.6 % (treated). For pou2, CpG1, 2, 3, 4, 5 are located respectively at the sites 96, 113, 178, 181, 212 downstream of the putative transcription start site (See Depince et al, 2021 for details). Their average methylation status was 30.8 ± 8.8 % (control) and 22.8 ± 3.0 % (treated). No significant effect of the treatment was observed on these CpG sites.

**Supplementary Methods:**

*Cell culture*

All cell culture procedure has been described in detail in Chenais 2019. The whole fin was minced and digested for 30 min with 2 mg/mL collagenase (Sigma C2674) in Leibovitz's L15 culture medium (Sigma L5520) supplemented with 5 mM Hepes, 2 mM $NaHCO_3$ 2 mM, 100 µg/mL gentamycin, 2.5 µg/mL amphotericin B (osmolality 290 mOsm/kg, pH 7.3) and 10% fetal calf serum, 2mM L-glutamine, 1% non-essential amino acids and 1% sodium pyruvate. Cells were plated in 6-well plates in growth L15 culture medium. After 24 hours, the supernatant enriched with mesenchymal cells, adhering more slowly than epithelial cells, was collected. These cells have previously been shown to be the most suitable for nuclear transfer (Chenais 2014, 2015). After filtration and washing, the cells derived from supernatant were seeded at $0.2.10^6$ cells in 24 well plates on 1.3 $cm^2$ glass coverslips and cultured in L15 medium for 2 days (about 80% confluence) until treatment experiments.

*RTqPCR analysis*

All steps of RT and qPCR were described in Chenais 2015. Briefly, 500 ng of total RNA was used to reverse transcription (RT) with the GoScript®™ Reverse Transcriptase System (A5001, Promega). Control reactions (RT- controls) were performed without the GoScript reverse transcriptase. All samples and controls were diluted 1/15 prior to qPCR. The primer sequences, concentrations and annealing temperatures used for col1a1a, nanog, pou2, sox2, c-myca1 and c-myca2, cDNA detection are described in Chenais 2015. The qPCR reactions were carried out in duplicates from 5 µL of cDNA samples or negative controls, 6 µL of the SYBR® Green Master Mix (Applied Biosystems) and 1 µL of reverse and forward primers mix. PCR were run on a StepOne real-Time PCR System (Applied Biosystems). Specificity of the PCR product was checked for each primer set and samples from the melting curve analysis.

Serial dilutions of fin or embryos cDNA (standard serial dilutions) depending on the target gene were run in duplicate for each gene in order to check the efficiency, linearity of the amplification for each gene and to determine the mean cycle threshold value (Ct) of the samples. The Ct values were normalized using the endogenous *18S rRNA* control and the target mRNA relative abundance was calculated according the formula: $2^{-\Delta Ct}$ with $\Delta Ct$ = mean Ct (*target gene*) – mean Ct (*18S rRNA*).

*Gene candidate DNA methylation analysis*

Total extracted DNA was purified using the Genomic DNA Purification and Concentration Kit (Zymo Research, D4010) and quantified using the QubitTM dsDNA HS Assay Kit (Q32851, Invitrogen) on the QubitTM 4 Fluorometer. DNA (10ng) was treated with bisulphite using the EZ DNA Methylation-Gold kit (Zymo Research, D5006). The bisulphited DNA was stored in a final volume of 12 µL at -20°C and analysed within 3 months of conversion. PCR reactions were performed to amplify the bisulphite sequences of regions of interest (promoter regions previously selected by the team), containing marker CpG sites, in each sample. Each reaction was performed in a final volume of 25 µL, using 2 µL of bisulphited DNA, the Advantage® 2 Polymerase enzyme (100X, Takara, 639202), the Advantage® 2 PCR buffer (10X, Takara, 639137), 0.2 µL of dNTPs (25 mM each, PROMEGA, U1330), 2 µL of primers (5 µM each, forward and reverse) and sterile water free of nucleases (PROMEGA, P1193). The reaction was carried out after denaturation of the components for 2 min at 94°C. The number of cycles as well as the temperature of hybridization of the primers is referenced in Depince et al., 2021. A second PCR serie (PCR nests) was carried out, using second pairs of primers selected from the ends of the amplicons generated by the first pairs of primers. Reverse primers of this nested PCR were coupled with biotin at their 5'end, in order to biotinylate the strands of amplified DNA, which is necessary for the subsequent sequencing steps. Each nested PCR reaction was performed in a final 50 µL volume, in duplicate, with 2 µL of products from the first PCR diluted 1/20th and following the same protocol as for the first reactions. Information about cycle number, hybridization temperature of the nested PCR primers is also given Depince et al., 2021. The nested PCR duplicates were then pooled and stored at -20°C. The quality of the nested PCR products (size and presence of a single amplicon) was tested by electrophoresis on a 2% agarose gel (Eurogenetec, EP-0010-05) including fluorescent nucleic acid intercalant (10,000X Red Nucleic Acid Gel Stain, Biotum).
